# *Bacteroidales* as a Fecal Contamination Indicator in Fresh Produce Industry: A Baseline Measurement

**DOI:** 10.1101/2023.07.17.549363

**Authors:** Jiangshan Wang, Mohsen Ranjbaran, Mohit S. Verma

**Affiliations:** Department of Agricultural and Biological Engineering, Purdue University, West Lafayette, IN 47906, USA; Birck Nanotechnology Center, Purdue University, West Lafayette, IN 47906, USA; Weldon School of Biomedical Engineering, Purdue University, West Lafayette, IN 47906, USA

**Keywords:** Fecal Contamination, *Bacteroidales*, Fresh Produce, baseline study

## Abstract

Foodborne outbreaks caused by fecal contamination of fresh produce represent a serious concern to public health and the economy. As the consumption of fresh produce increases, public health officials and organizations have pushed for improvements in food safety procedures and environmental assessments to reduce the risk of contamination. Visual inspections and the establishment of “buffer zones” between animal feeding operations and producing fields are the current best practices for environmental assessments. However, a generalized distance guideline and visual inspections may not be enough to account for all environmental risk variables. Here, we report a baseline measurement surveying the background *Bacteroidales* concentration, as a quantitative fecal contamination indicator, in California’s Salinas Valley. We collected a total of 1632 samples from two romaine lettuce commercial fields at the time of harvesting through two seasons in a year. The *Bacteroidales* concentration was very low (0 – 2.00 copies/cm^2^). Furthermore, we established a practical methodology for evaluating the risk of fecal contamination in a real-world setting, complementing the current environmental assessment practices. This method can identify site-specific risks and offer fresh produce stakeholders a more comprehensive understanding of their fields. We anticipate this work can encourage the use of *Bacteroidales* in the fresh produce industry to monitor fecal contamination and prevent future foodborne outbreaks.

## 1. Introduction

Fresh produce can serve as a vehicle for foodborne pathogens and has been associated with a significant number of multistate outbreaks in the United States over the years (CDC, 2022; Pang et al., 2018). Fresh produce is generally cultivated in open fields, making it susceptible to environmental reservoirs of foodborne pathogens (such as poorly composted animal manures, subpar irrigation water, encroachment of wild animals, and bioaerosols from nearby animal operations) during production (Alegbeleye et al., 2018; Chen et al., 2017; Li et al., 2023).

Many fresh produce organizations have devised and implemented safety practices and protocols to reduce potential sources of contamination. The California Leafy Greens Marketing Agreement (LGMA) Food Safety Standards, for example, specify that the best practice for environmental assessments is to inspect the production field and surrounding area for potential animal hazards or other sources of human pathogens of concern. A “buffer zone” of 400 feet for animal feeding operations (less than 1,000 animals) or 1200 feet for concentrated animal feeding operations (1,000-80,000 animals) around the production field is required to prevent pathogen transmission from animals to crops (LGMA, 2021). However, the fact that each farm has a unique combination of environmental risk variables (e.g., topography, land-use interactions, and weather) makes this generalized distance guideline difficult to justify (Strawn et al., 2013a, 2013b). Furthermore, while these practices were initially used to limit food safety risks for preharvest production, applying them to all fields without specificity would elevate production costs (for low-risk fields) and raise produce safety concerns (for high-risk fields). Indeed, LGMA acknowledges that there is limited information on which to base this recommendation, and ideally an appropriate “buffer zone” should be customized to each farm (Hoar, 2011; Strawn et al., 2013b).

Fecal indicator bacteria (FIB) such as *Escherichia coli*, *Enterococcus faecalis,* and *Bacteroidales,* have been used to assess possible fecal contamination in fresh produce (Denis et al., 2016; Drozd et al., 2013; Harris et al., 2017; Ordaz et al., 2019). Certain features of *Bacteroidales* make them superior to other FIB. These features include high prevalence in feces (constituting 30%–40% of total fecal bacteria, 10^9^ to 10^11^ colony forming units (CFU)/g), obligate anaerobicity (preventing their growth and multiplication in the ambient environment), low natural abundance from non-fecal sources, and high host-specificity (various sequences of the 16S rRNA gene have been designed to detect fecal pollution from specific hosts) (Mascorro et al., 2018; Ordaz et al., 2019). As a result, *Bacteroidales* serve as a valuable—and potentially quantitative—marker in each farm, not only to assess the risk of contamination based on the farm’s unique combination of environmental risk variables but also to track the source and resolve fecal contamination. However, the levels of *Bacteroidales* naturally present in the environment of various fresh produce operations remain undetermined so far.

In our previous study, we developed a *Bacteroidales* detection assay and combined it with a new method for collecting bioaerosols to perform a field-deployable risk assessment of fecal contamination (Wang et al., 2023). We also quantified the concentration of *Bacteroidales* in fields adjacent to animal feeding operations using a conventional quantitative polymerase chain reaction (qPCR) assay and associated that concentration of *Bacteroidales* with a high risk of fecal contamination (Wang et al., 2023). Here, we have surveyed the baseline concentration of *Bacteroidales* in commercial fields committed to safe production standards. This baseline study has significant practical implications for fecal contamination management in the fresh produce industry, including i) interpreting the possibility of fecal contamination from *Bacteroidales* concentrations, ii) implementing the developed *Bacteroidales* detection assay (Wang et al., 2023), as well as other *Bacteroidales* assays in the fresh produce industry, and iii) assisting fresh produce growers to determine site-specific risk and their decision-making process regarding the microbial safety of fresh produce. Although *Bacteroidales* do not naturally occur in the environment, a very low amount of *Bacteroidales* DNA and genetic material of other fecal indicator bacteria could be present in the production field via several routes (e.g., organic fertilizer, residuals from previous contamination, etc.).

Therefore, to apply *Bacteroidales* for monitoring fecal contamination in fresh produce production, a baseline study needs to be conducted to survey the background concentration of *Bacteroidales* in “low-risk” fresh produce fields. Owing to support from the fresh produce industry, our research team was provided a unique opportunity to survey fecal contamination in fresh produce under real-world conditions. In this study, we collected a total of 1632 samples from two romaine lettuce commercial fields in California’s Salinas Valley at the time of harvesting over two seasons in a year. We hope to establish the baseline measurement of *Bacteroidales* in fields with low fecal contamination risk and support the use of *Bacteroidales* in the fresh produce industry for assessing the risk of both general fecal contamination and microbial source tracking.

## 2. Materials and Methods

### 2.1 Sample collection

Samples were collected between May 2021 and August 2021 over two growing seasons in two romaine lettuce commercial fields in Salinas, California, United States. The fields were labeled with row and column numbers with the distance between each row and column to be 6 meters. Samples were collected at the intersection of each row and column (approximately 100 sampling sites per acre of field). Two types of samples were collected at each sampling site 1) 25 g of romaine lettuce leaf sample (approximately four leaves) 2) collection flag sample. The sample size for the romaine lettuce leaf sample was determined following United States Food and Drug Administration (FDA) Bacteriological Analytical Manual (BAM) for isolating specific pathogens from fresh vegetable samples (FDA, 2021). The fabrication and deploying method for the collection flags are described elsewhere (Wang et al., 2023). Briefly, the collection flags were assembled with a bamboo skewer (29.8 cm) and a piece of transparent film pre-cut into 7.62 × 21.59 cm (3 × 8.5 inches) strips. One collection flag was placed at each sampling site 7 days before the sample collection.

During the first growing season (May 2021), we collected 336 romaine lettuce leaf samples and 336 collection flag samples over a field size of 3.3 acres (16 rows, 21 columns) (Figure 1A). In the second growing season (August 2021), we collected 480 romaine lettuce leaf samples and 480 collection flag samples over a field size of 4.8 acres (8 rows, 60 columns) (Figure 2A). We collected the samples wearing a Tyvek suit, gloves, and a mask to avoid contaminating the samples (Figure 1). Before collecting each sample, we used 70% ethanol to sanitize gloves and sleeves to avoid cross-contamination. Each sample was placed in an individual, pre-labeled Ziploc resealable storage bag (B07NQVYCG3, Amazon, USA). The collected samples were kept on ice and shipped back to West Lafayette via FedEx Priority Overnight in a cooler box with ice packs. Following sample collection, the remaining lettuce in the experimental fields was destroyed in the field.

**Figure 1:**
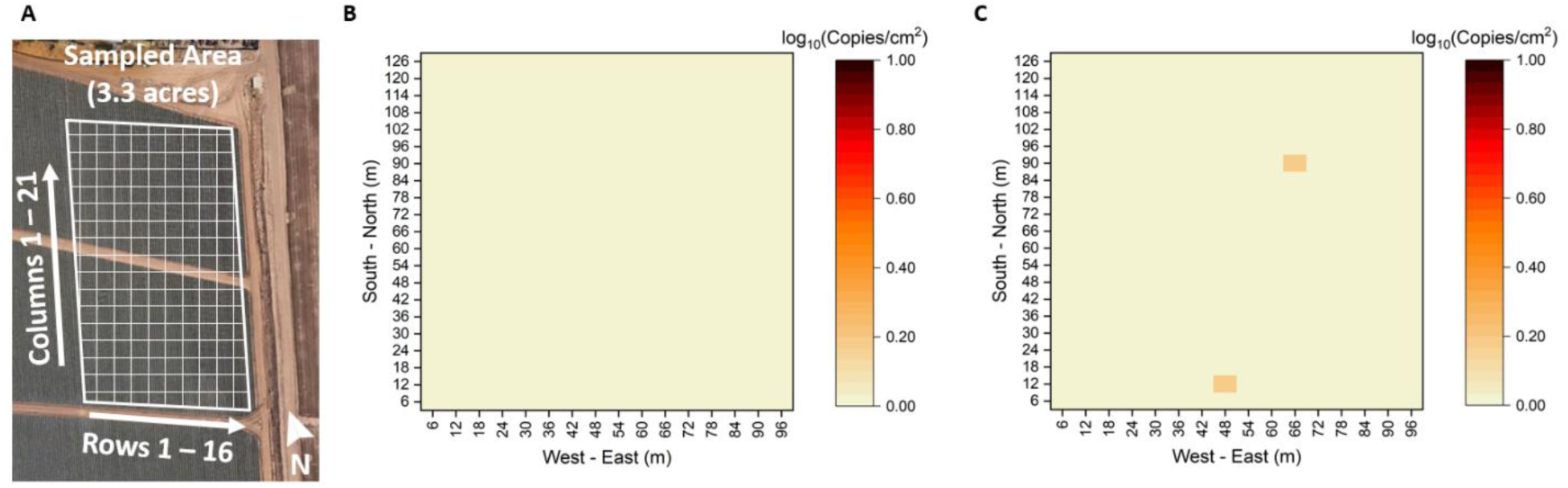
Risk of fecal contamination mapping using qPCR (May 2021). The Ct value of each qPCR reaction was converted to log_10_ (copies/cm^2^) via a linear fit to log-transformed concentrations.

**Figure 2:**
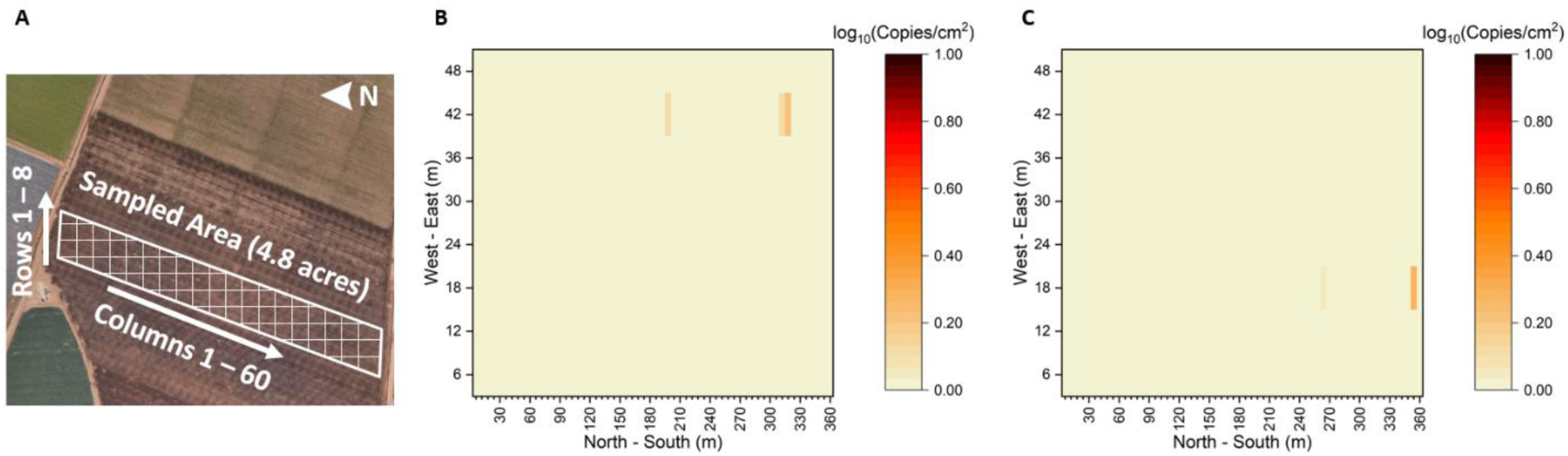
Risk of fecal contamination mapping using qPCR (August 2021). The Ct value of each qPCR reaction was converted to log_10_ (copies/cm^2^) via a linear fit to log-transformed concentrations.

### 2.2 Sample processing

The samples were stored at 4°C after receiving them. The romaine lettuce leaf samples were processed by a washing and filtering method modified from the FDA BAM for isolating specific pathogens from fresh vegetable samples (FDA, 2021). 225 mL of ultrapure water (PURELAB flex, ELGA, USA) was added to the sealed bag (B07NQVYCG3, Amazon, USA) with 25 g of lettuce leaf sample. The bag was hand-shaken for 1 min to elute any bacteria into the solution. The wash solution was then filtered using a 90 mm, 0.22 µm, cellulose acetate (CA) membrane (FBM090CA022, Filter-Bio, China). The filtered membrane was immersed into 1 mL of nuclease-free water inside a 2 mL centrifuge tube and the tube was vortexed at maximum speed for 1 min. Finally, the tube was centrifuged at 10,000 rpm for 1 min to recover the resuspension. The membrane was removed from the tube after centrifugation. Each collection flag was swabbed using a wet polyester-tipped swab (263000, BD BBL, USA) and was resuspended in 200 μL nuclease-free water. All samples were kept at -20 °C until the experiment.

### 2.3 Genomic DNA preparation

*B. fragilis* (ATCC^®^ 25285^™^) was grown overnight (37°C, 4% H_2_, 5% CO_2_, 91% N_2_, <10 ppm O_2_) in Chopped Meat Carbohydrate Broth (BD297307, BD, USA). Genomic DNA was extracted from *B. fragilis* with Purelink Genomic DNA Mini Kit (K182001, Invitrogen, USA) according to the manufacturer’s protocol. The extracted DNA product was quantified using Quant-iT™ PicoGreen™ dsDNA Assay Kit (P7589, Thermo Fisher, USA).

### 2.4 Quantitative PCR (qPCR)

The qPCR reactions were performed in a total volume of 20 μL, containing 10 μL 2X Luna® Universal Probe qPCR Master Mix (M3004, New England Biolabs, USA) (final concentration 1X), 0.8 μL of 10 μM forward primer (final concentration 0.4 μM), 0.8 μL of 10 μM reverse primer (final concentration 0.4 μM), 0.4 μL of 10 μM fluorescent probe (final concentration 0.2 μM) (Table 1), 7 μL nuclease-free water, and 1 μL of template or 1 μL of nuclease-free water for no template control (NTC). The resuspensions of both membrane and swab were directly used for qPCR assays without performing DNA extraction. The qPCR reactions were performed on a qTOWER3 Real-Time Thermal Cycler (Analytik Jena, Germany), and the thermal cycling conditions were implemented using the following program: initial denaturation at 95 °C for 1 min, followed by 45 cycles of 95 °C for 15 s, 55°C for 15 s, and 60 °C for 30 s.

**Table 1:**
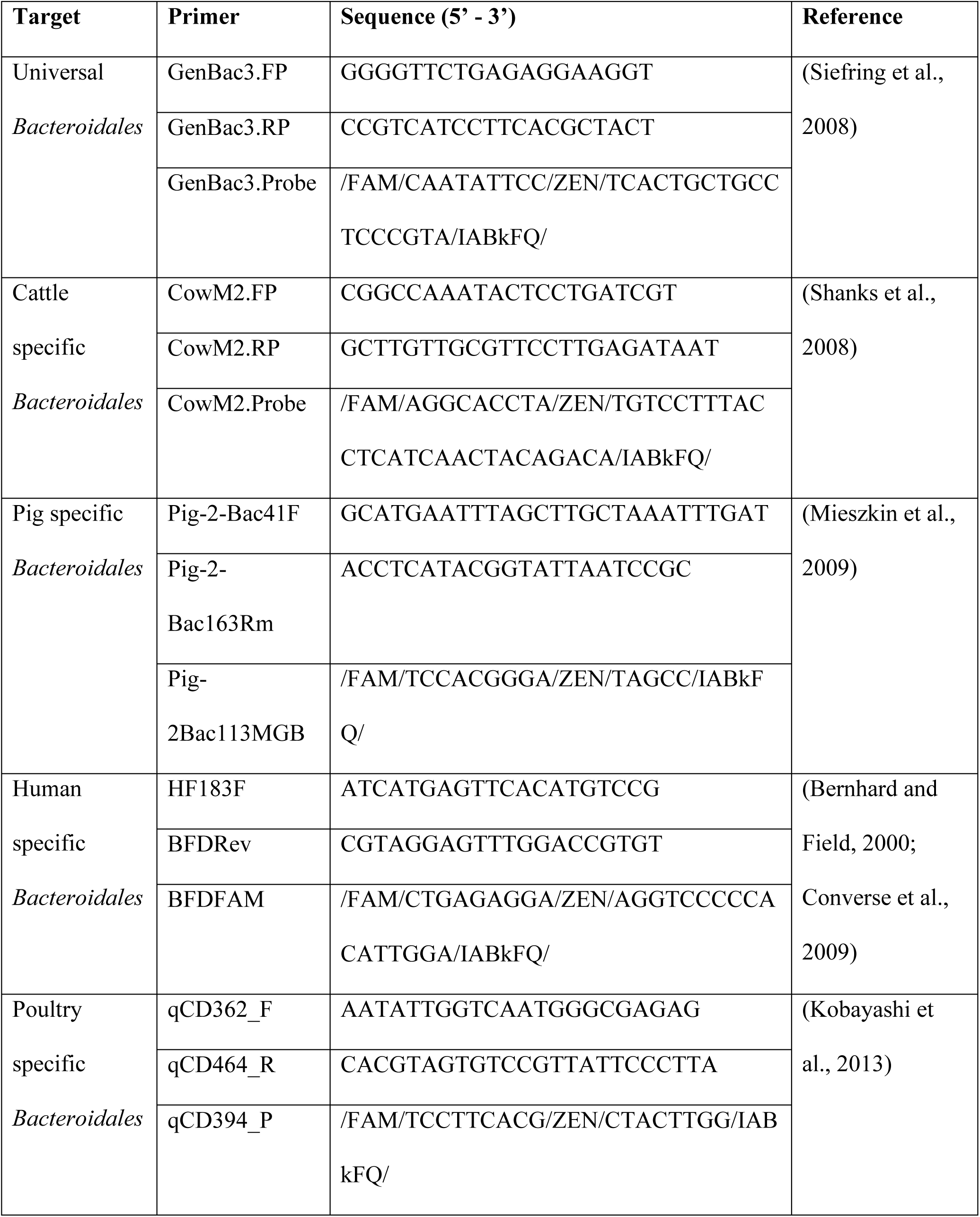
Sequences for primers and probes used in this study.

### 2.5 Digital PCR (dPCR)

The dPCR reactions were performed in a total volume of 12 μL, containing 3 μL 4X Probe PCR Master Mix (250102, Qiagen, USA) (final concentration 1X), 1.2 μL of 10X primer-probe mix (final concentration 1X, 0.8 μM forward primer, 0.8 μM reverse primer, 0.4 μM FAM probe), 2.8 μL nuclease-free water, and 5 μL of the template or 5 μL of nuclease-free water for NTC. 10X primer-probe mix is one of the host-specific qPCR primer-probe set in Table 1 (cattle-specific *Bacteroidales*, swine-specific *Bacteroidales*, human-specific *Bacteroidales*, and poultry-specific *Bacteroidales*). The dPCR reactions were performed in an 8.5K 96-well Nanoplate (250021, Qiagen, USA) on a 5-plex QIAcuity One digital PCR instrument (911021, Qiagen, USA). The thermal cycling conditions were implemented using the following program: initial denaturation at 95 °C for 2 min, followed by 40 cycles of 95 °C for 15 s, 55 °C for 15 s, and 60 °C for 30 s.

## 3. Results

### 3.1 Construction of qPCR calibration curve

The fluorescence intensities were extracted for the 45-min time point for qPCR reactions. Any fluorescent intensity values that were greater than the highest background intensity (20% of the maximum reaction intensity) were considered successful amplifications. The lowest DNA concentration that had successful amplification for all three replicates was classified as the limit of detection (LoD). The qPCR showed an LoD of 1 copy/reaction. The Ct values (number of cycles required for fluorescent intensity to reach/exceed defined reaction threshold) for qPCR were calculated using software qPCRsoft 4.1 (baseline correction: 5, auto threshold) (Analytik Jena, Germany), and reported in Table 2. Linear regression analysis was used to fit correlations between Ct values and log_10_(copies/reaction) (Figure S1).

**Table 2:** Limit of detection characterization of the assay. *B. fragilis* genomic DNA (1 μL) was added to reactions (19 μL reagents) in triplicate at different concentrations (serially diluted from 1000 copies/reaction to 1 copy/reaction). The Ct value (in minutes) of each reaction is reported in the table.

| DNA concentration<br>(copies/reaction) | *Bacteroidales* |  |  |
| --- | --- | --- | --- |
|  | qPCR (GenBac3) |  |  |
| 1000 | 24.12 | 24.10 | 24.04 |
| 500 | 24.73 | 25.02 | 25.03 |
| 100 | 27.33 | 27.33 | 27.34 |
| 50 | 28.34 | 28.89 | 28.48 |
| 25 | 29.66 | 29.71 | 29.08 |
| 10 | 31.21 | 30.74 | 31.33 |
| 5 | 31.87 | 32.39 | 32.34 |
| 1 | 34.75 | 35.95 | 35.89 |
| 0 | No Ct | No Ct | No Ct |

### 3.2 *Bacteroidales* concentration on field-grown lettuce and collection flags

The processed samples were used for qPCR assays. The fluorescence intensities were extracted for the 45-min time point for qPCR reactions. The Ct values for qPCR were calculated using the software qPCRsoft 4.1 (baseline correction: 5, auto threshold) (Analytik Jena, Germany). The Ct value of each sample was then used to calculate the *Bacteroidales* concentration using the constructed calibration curve. To construct a fecal contamination risk evaluation map, the *Bacteroidales* concentration result of both types of samples (romaine lettuce leaf samples and collection flag samples) were normalized to log_10_(copies/cm^2^) (Figure 1 and Figure 2).

During May 2021 harvesting season, we collected 672 samples (including 336 lettuce leaf samples and 336 collection flag samples) from a sampled area of 3.3 acres. For most samples, *Bacteroidales* was not detected via qPCR, only 9 samples (leaf samples: R3C19, R11C16; flag samples: R2C20, R4C17, R6C1, R8C2, R10C2, R11C15, R16C6; R: row, C: column) returned a *Bacteroidales* concentration higher than 1 copy/reaction, which is the qPCR assay’s limit of detection. The *Bacteroidales* concentrations of these samples are reported in Table 3. During the August 2021 harvesting season, we collected 960 samples (including 480 lettuce leaf samples and 480 collection flag samples) from a sampled area of 4.8 acres. Similar to the result from samples collected in May 2021, only 8 samples (leaf samples: R6C32, R7C33, R7C52, R7C53; flag samples: R3C44, R3C59, R5C1, R8C11; R: row, C: column) returned a *Bacteroidales* concentration higher than 1 copy/reaction (Table 3). As expected, the concentrations of *Bacteroidales* in both romaine lettuce commercial fields from two harvesting seasons were very low (0 – 2.00 copies/cm^2^).

**Table 3:** Samples selected for microbial source tracking.

| Collection time | Sample type | Sample ID | Copies/cm<sup>2</sup> |
| --- | --- | --- | --- |
| May 2021 | Leaf sample | R3C19 | 0.88 |
|  |  | R11C16 | 0.87 |
|  | Collection flag | R2C20 | 0.62 |
|  |  | R4C17 | 0.98 |
|  |  | R6C1 | 0.62 |
|  |  | R8C2 | 1.58 |
|  |  | R10C2 | 0.82 |
|  |  | R11C15 | 1.57 |
|  |  | R16C6 | 0.69 |
| August 2021 | Leaf sample | R6C32 | 0.82 |
|  |  | R7C33 | 1.32 |
|  |  | R7C52 | 1.28 |
|  |  | R7C53 | 1.67 |
|  | Collection flag | R3C44 | 1.11 |
|  |  | R3C59 | 2.00 |
|  |  | R5C51 | 0.63 |
|  |  | R8C11 | 0.94 |

### 3.3 Microbial source tracking

Samples that returned a concentration higher than 1 copy/reaction (17 samples) were selected for microbial source tracking. Each sample was tested with four different host-specific qPCR primer-probe sets (cattle-specific *Bacteroidales*, swine-specific *Bacteroidales*, human-specific *Bacteroidales*, and poultry-specific *Bacteroidales*) (Table 1).

Since the host-specific populations represent a small group within the general *Bacteroidales* population (Bernhard and Field, 2000; Lamendella et al., 2007), a dPCR method was adopted for the microbial source tracking experiment to achieve a higher level of sensitivity (Milbury et al., 2014). dPCR is commonly used in environmental research (Amoah et al., 2022; Guérin-Rechdaoui et al., 2022; Rumky et al., 2022) and due to the inherent nature of dPCR, the assay has a high tolerance to biological inhibitors and has better performance on trace detection for a minority target (Milbury et al., 2014; Perkins et al., 2017).

Following plate preparation, the system partitioned each sample into approximately 8500 partitions, with approximately 8300 valid counts. Each partition was individually sealed following 40 cycles of thermocycling. The plate was then imaged to count the number of positive/fluorescent partitions for each sample. The fluorescent threshold was determined to be 20 relative fluorescence units (RFU) based on the NTC. 16 partitions were counted as positive, including 2 positive partitions for cattle-specific *Bacteroidales*, 3 positive partitions for swine-specific *Bacteroidales*, 2 positive partitions for human-specific *Bacteroidales*, and 9 positive partitions for poultry-specific *Bacteroidales* (Figure 3). We were unable to make definitive statements about microbial source tracking due to the low copy number of host-specific *Bacteroidales*.

**Figure 3:**
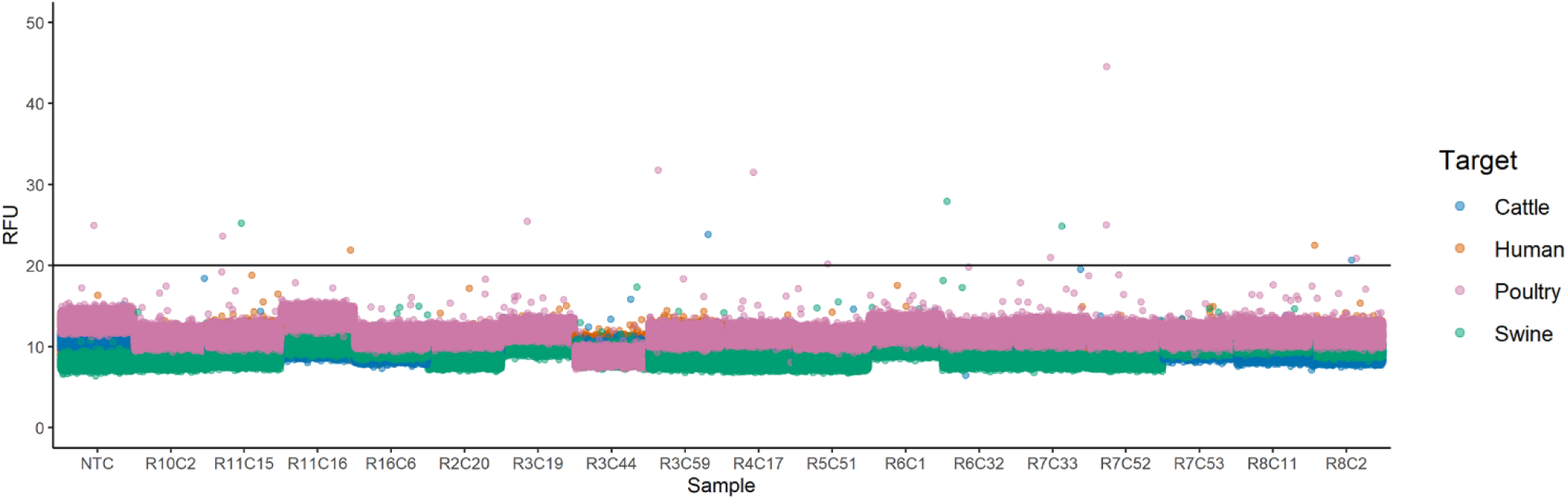
Scatter plot of microbial source tracking result. Each of the 17 samples with a concentration greater than 1 copy/reaction was selected for microbial source tracking. The experiment was carried out in a dPCR method. Each sample was tested with four different host-specific qPCR primer-probe sets (cattle-specific *Bacteroidales*, swine-specific *Bacteroidales*, human-specific *Bacteroidales*, and poultry-specific *Bacteroidales*). On the scatter plot, the fluorescence intensity of each partition was displayed. Based on the NTC, the fluorescence threshold was determined to be 20 RFU.

## 4. Discussion

### 4.1 *Bacteroidales* concentration as a fecal contamination biomarker in fresh produce fields

*Bacteroidales* have been extensively employed as a water fecal contamination indicator (Cyterski et al., 2022; Ekhlas et al., 2021; Nshimyimana et al., 2017). Few studies have used *Bacteroidales* to evaluate potential fecal contamination in fresh produce (Mascorro et al., 2018; Ordaz et al., 2019; Ravaliya et al., 2014; Wang et al., 2023). Advantages of implementing *Bacteroidales* include: i) high abundance in contaminated samples and feces; ii) obligate anaerobicity, preventing their growth and multiplication in the ambient environment; iii) low natural abundance from non-fecal sources; and iv) high host-specificity, making it suitable for microbial source tracking – all of which making it a valuable marker for detecting fecal contamination in the fresh produce industry.

To support the use of *Bacteroidales* for monitoring fecal contamination in fresh produce production, in the present study, we determined the baseline concentration of *Bacteroidales* under real-world conditions in two romaine lettuce commercial fields in California’s Salinas Valley at the time of harvesting over two seasons in a year. Both production fields comply with safe production standards, therefore, this baseline study reflects the *Bacteroidales* level in fresh produce fields with “low risk” fecal contamination. We collected a total of 1632 samples consisting of two types of samples (romaine lettuce leaf samples and collection flag samples) from the field and processed each type of sample with a different method (a conventional washing and filtering method modified from FDA BAM for lettuce leaf samples and a swabbing method modified from our previous work for collection flag samples) (FDA, 2021; Wang et al., 2023). Among the 1632 samples, only 17 samples returned a concentration higher than 1 copy/reaction (∼ 1%). This finding is consistent with the hypothesis that, while *Bacteroidales* do not normally occur in the environment, a trace amount of *Bacteroidales* DNA may be present in the field as a result of various reasons (e.g., organic fertilizer, residuals from previous contamination, etc.).

In our previous study, we determined the concentration of *Bacteroidales* in fields with “high risk” fecal contamination (adjacent to animal feeding operations) (Wang et al., 2023). The concentration was quantified at over 10^4^ copies/cm^2^ (of *Bacteroidales* per cm^2^), compared to the highest concentration observed in this study of 2 copies/cm^2^. According to the observations, there is a significant difference of about four orders of magnitude in the concentration of *Bacteroidales* between “high risk” and “low risk” of fecal contamination. This distinction could be exploited to detect fecal contamination events in the fields; for instance, a hotspot on heatmaps (as shown in Figures 1 and 2) could indicate spot contaminations such as wild animal intrusion. A “gradient of contamination” could be seen for continuous contaminations (animal feeding operations, irrigation water), indicating the source of contamination. Such outcomes are part of ongoing work in our lab and will be published in the future.

Although we intended to assess contamination levels between different seasons (whether a season is more susceptible to fecal contamination) and whether one collecting/processing method is preferable to the other, at a very low concentration, we do not see a significant difference between seasons or different collecting/processing methods. In terms of applicability, however, collection flags offer an advantage over lettuce leaf samples by providing a standalone carrier for measuring fecal contamination. Environmental assessments are required at several phases throughout the production cycle, including before the vegetation planting. Growers must identify any potential sources of contamination in the production field and determine an adequate “buffer zone” to minimize environmental risks during production. The collecting/processing method employing collection flags allows growers to conduct field assessments even without a product in the field.

The presence of *Bacteroidales* indicates fecal contamination, however, fecal contamination is not always associated with the presence of enteric pathogens. FIBs are normally present in much higher concentrations than any of the pathogens and are also more constantly detected in stool samples, compared to pathogens (Korajkic et al., 2018). As a result, focusing exclusively on pathogen screening could deliver a false-negative result and conceal the fact that there is a high risk of fecal exposure in the field. If an extraordinarily high level of *Bacteroidales* was detected in the field, regardless of the presence or absence of pathogens, it implies that the field has been exposed to serious fecal contamination, and the grower must act immediately to remedy the exposure. Additionally, as visual inspection is the only gold standard test currently available for field assessments, we intend to offer fresh produce stakeholders an alternative by providing a quantitative understanding of the potential of fecal contamination in their fields. In the future, it is also possible to conduct these risk assessments in the field by using assays such as loop-mediated isothermal amplification (LAMP) (Davidson et al., 2021; Mohan et al., 2021; Pascual-Garrigos et al., 2021; Ranjbaran and Verma, 2022; Wang et al., 2023, 2022, 2021) as opposed to lab-based qPCR.

### 4.2 Using *Bacteroidales* for microbial source tracking

Host-specific *Bacteroidales* markers are usually evaluated with fecal samples or highly contaminated environment samples (e.g., sewages, wastewater treatment plants, and water bodies close to animal feeding operations) (Fremaux et al., 2009; Malla et al., 2018). These markers were selected to be strictly host-specific, thus there shouldn’t be any cross-amplification with fecal DNA from other species. As a result, these host-specific *Bacteroidales* constitute a relatively small proportion of the total *Bacteroidales* population and are thus not always present when fecal contamination occurs, particularly at low levels of contamination (Pendergraph et al., 2021; Silkie and Nelson, 2009). When choosing a host-specific marker, there is a trade-off between specificity and sensitivity; where a decreased cross-reaction rate between various hosts might also affect the sensitivity of the assay. Therefore, microbial source tracking using host-specific *Bacteroidales* markers may not be appropriate for low-level fecal contamination fields. Furthermore, for studies attempting to detect fecal contamination, we strongly recommend using not only host-specific *Bacteroidales* markers but also a universal *Bacteroidales* marker to reduce the incidence of false negatives.

## 5. Conclusions

In conclusion, this work is the first reported baseline study surveying the background concentration of *Bacteroidales* as a fecal contamination indicator in fresh produce fields. We were able to survey fecal contamination in fresh produce fields under real-world circumstances in Salinas, one of the major fresh-produce producing areas in the USA. As expected, the concentrations of *Bacteroidales* in the sampled commercial fields were very low (0 – 2.00 copies/cm^2^). On the other hand, “high risk” fecal contamination (adjacent to animal feeding operations) areas are reported to have *Bacteroidales* concentrations over 10^4^ copies/cm^2^ (Wang et al., 2023). There is a significant difference in the concentration of *Bacteroidales* between “high risk” and “low risk” of fecal contamination. Therefore, *Bacteroidales* can be a valuable tool for researchers undertaking comprehensive studies to monitor fecal pollution and facilitate microbial safety in the fresh produce industry. We attempted to perform microbial source tracking with host-specific *Bacteroidales* markers, however, due to the low concentration of host-specific *Bacteroidales*, we were unable to draw definitive statements about the source of fecal contamination.

In addition, we demonstrated a practical methodology for evaluating the risk of fecal contamination in a real-world setting. As a complementary practice to the current environmental assessments (via visual inspections), we proposed a more quantitative solution for fresh produce stakeholders to have a more comprehensive understanding of the potential of fecal contamination in their fields. This method can determine site-specific risk and determine an adequate “buffer zone” to minimize environmental risks, thereby avoiding unnecessary production costs caused by unified safety practices.

We hope that this baseline *Bacteroidales* measurement in fields with low fecal contamination risk can support the use of *Bacteroidales* in the fresh produce industry for assessing the risk of fecal contamination and preventing future foodborne disease outbreaks.

## Author contributions

**Jiangshan Wang:** Conceptualization, Investigation, Data Curation, Visualization, Writing – Original Draft, Writing - Review & Editing. **Mohsen Ranjbaran:** Conceptualization, Investigation, Writing - Review & Editing. **Mohit S. Verma:** Conceptualization, Supervision, Writing – Review & Editing, Funding acquisition

## Acknowledgment

This work was funded by the Center for Produce Safety (CPS Award Number: 2021CPS12), the California Department of Food and Agriculture (CDFA Agreement No. 20-0001-054-SF), and the U.S. Department of Agriculture’s (USDA) Agricultural Marketing Service (USDA Cooperative Agreement No. USDA-AMS-TM-SCBGP-G-20-0003). Any opinions, findings, conclusions, or recommendations expressed in this publication or audiovisual are those of the author(s) and do not necessarily reflect the views of The Center for Produce Safety, the California Department of Food and Agriculture, or the U.S. Department of Agriculture’s (USDA) Agricultural Marketing Service. The project entitled “Field evaluation of microfluidic paper-based analytical devices for microbial source tracking” was funded in whole or in part through a subrecipient grant awarded The Center for Produce Safety through the California Department of Food and Agriculture 2020 Specialty Crop Block Grant Program and the U.S. Department of Agriculture’s (USDA) Agricultural Marketing Service.

## Data Availability

The datasets generated during and/or analyzed during the current study are available in the Mendeley Data repository, doi: 10.17632/gsxhh3f4cr.1.

## References

1. Alegbeleye, O.O., Singleton, I., Sant’Ana, A.S., 2018. Sources and contamination routes of microbial pathogens to fresh produce during field cultivation: A review. Food Microbiol. 73, 177–208. https://doi.org/10.1016/j.fm.2018.01.003

2. Amoah, I.D., Abunama, T., Awolusi, O.O., Pillay, L., Pillay, K., Kumari, S., Bux, F., 2022. Effect of selected wastewater characteristics on estimation of SARS-CoV-2 viral load in wastewater. Environ. Res. 203, 111877. https://doi.org/10.1016/j.envres.2021.111877

3. Bernhard, A.E., Field, K.G., 2000. A PCR Assay To Discriminate Human and Ruminant Feces on the Basis of Host Differences in Bacteroides-Prevotella Genes Encoding 16S rRNA. Appl. Environ. Microbiol. 66, 4571–4574. https://doi.org/10.1128/AEM.66.10.4571-4574.2000

4. CDC, 2022. Summary of Possible Multistate Enteric (Intestinal) Disease Outbreaks in 2017– 2020 [WWW Document]. URL https://www.cdc.gov/foodsafety/outbreaks/multistate-outbreaks/annual-summaries/annual-summaries-2017-2020.html (accessed 12.5.22).

5. Chen, J.-Q., Regan, P., Laksanalamai, P., Healey, S., Hu, Z., 2017. Prevalence and methodologies for detection, characterization and subtyping of Listeria monocytogenes and L. ivanovii in foods and environmental sources. Food Sci. Hum. Wellness 6, 97–120. https://doi.org/10.1016/j.fshw.2017.06.002

6. Converse, R.R., Blackwood, A.D., Kirs, M., Griffith, J.F., Noble, R.T., 2009. Rapid QPCR-based assay for fecal Bacteroides spp. as a tool for assessing fecal contamination in recreational waters. Water Res., Cross-validation of detection methods for pathogens and fecal indicators 43, 4828–4837. https://doi.org/10.1016/j.watres.2009.06.036

7. Cyterski, M., Shanks, O.C., Wanjugi, P., McMinn, B., Korajkic, A., Oshima, K., Haugland, R., 2022. Bacterial and viral fecal indicator predictive modeling at three Great Lakes recreational beach sites. Water Res. 223, 118970. https://doi.org/10.1016/j.watres.2022.118970

8. Davidson, J.L., Wang, J., Maruthamuthu, M.K., Dextre, A., Pascual-Garrigos, A., Mohan, S., Putikam, S.V.S., Osman, F.O.I., McChesney, D., Seville, J., Verma, M.S., 2021. A paper-based colorimetric molecular test for SARS-CoV-2 in saliva. Biosens. Bioelectron. X 9, 100076. https://doi.org/10.1016/j.biosx.2021.100076

9. Denis, N., Zhang, H., Leroux, A., Trudel, R., Bietlot, H., 2016. Prevalence and trends of bacterial contamination in fresh fruits and vegetables sold at retail in Canada. Food Control 67, 225–234. https://doi.org/10.1016/j.foodcont.2016.02.047

10. Drozd, M., Merrick, N.N., Sanad, Y.M., Dick, L.K., Dick, W.A., Rajashekara, G., 2013. Evaluating the Occurrence of Host-Specific Bacteroidales, General Fecal Indicators, and Bacterial Pathogens in a Mixed-Use Watershed. J. Environ. Qual. 42, 713–725. https://doi.org/10.2134/jeq2012.0359

11. Ekhlas, D., Kurisu, F., Kasuga, I., Cernava, T., Berg, G., Liu, M., Furumai, H., 2021. Identification of new eligible indicator organisms for combined sewer overflow via 16S rRNA gene amplicon sequencing in Kanda River, Tokyo. J. Environ. Manage. 284, 112059. https://doi.org/10.1016/j.jenvman.2021.112059

12. FDA, 2021. Bacteriological Analytical Manual (BAM).

13. Fremaux, B., Gritzfeld, J., Boa, T., Yost, C.K., 2009. Evaluation of host-specific Bacteroidales 16S rRNA gene markers as a complementary tool for detecting fecal pollution in a prairie watershed. Water Res., Cross-validation of detection methods for pathogens and fecal indicators 43, 4838–4849. https://doi.org/10.1016/j.watres.2009.06.045

14. Guérin-Rechdaoui, S., Bize, A., Levesque-Ninio, C., Janvier, A., Lacroix, C., Le Brizoual, F., Barbier, J., Amsaleg, C.R., Azimi, S., Rocher, V., 2022. Fate of SARS-CoV-2 coronavirus in wastewater treatment sludge during storage and thermophilic anaerobic digestion. Environ. Res. 214, 114057. https://doi.org/10.1016/j.envres.2022.114057

15. Harris, A.R., Islam, M.A., Unicomb, L., Boehm, A.B., Luby, S., Davis, J., Pickering, A.J., 2017. Fecal Contamination on Produce from Wholesale and Retail Food Markets in Dhaka, Bangladesh. Am. J. Trop. Med. Hyg. 98, 287–294. https://doi.org/10.4269/ajtmh.17-0255

16. Hoar, B.R., 2011. Developing buffer zone distances between sheep grazing operations and vegetable crops to maximize food safety. [WWW Document]. URL https://www.centerforproducesafety.org/researchproject/318/awards/Developing_buffer_zone_distances_between_sheep_grazing_operations_and_vegetable_crops_to_maximize_food_safety.html (accessed 12.5.22).

17. Kobayashi, A., Sano, D., Okabe, S., 2013. Effects of temperature and predator on the persistence of host-specific Bacteroides-Prevotella genetic markers in water. Water Sci. Technol. 67, 838–845. https://doi.org/10.2166/wst.2012.626

18. Korajkic, A., McMinn, B.R., Harwood, V.J., 2018. Relationships between Microbial Indicators and Pathogens in Recreational Water Settings. Int. J. Environ. Res. Public. Health 15, 2842. https://doi.org/10.3390/ijerph15122842

19. Lamendella, R., Domingo, J.W.S., Oerther, D.B., Vogel, J.R., Stoeckel, D.M., 2007. Assessment of fecal pollution sources in a small northern-plains watershed using PCR and phylogenetic analyses of Bacteroidetes 16S rRNA gene. FEMS Microbiol. Ecol. 59, 651–660. https://doi.org/10.1111/j.1574-6941.2006.00211.x

20. LGMA, 2021. Food Safety Program [WWW Document]. Calif. Leafy Greens Mark. Agreem. URL http://lgma.ca.gov/food-safety-program (accessed 12.5.22).

21. Li, B., Wang, H., Xu, J., Qu, W., Yao, L., Yao, B., Yan, C., Chen, W., 2023. Filtration assisted pretreatment for rapid enrichment and accurate detection of Salmonella in vegetables. Food Sci. Hum. Wellness 12, 1167–1173. https://doi.org/10.1016/j.fshw.2022.10.042

22. Malla, B., Ghaju Shrestha, R., Tandukar, S., Bhandari, D., Inoue, D., Sei, K., Tanaka, Y., Sherchand, J. b., Haramoto, E., 2018. Validation of host-specific Bacteroidales quantitative PCR assays and their application to microbial source tracking of drinking water sources in the Kathmandu Valley, Nepal. J. Appl. Microbiol. 125, 609–619. https://doi.org/10.1111/jam.13884

23. Mascorro, J.A., Hernández-Rangel, L.G., Heredia, N., García, S., 2018. Bacteroidales as Indicators and Source Trackers of Fecal Contamination in Tomatoes and Strawberries. J. Food Prot. 81, 1439–1444. https://doi.org/10.4315/0362-028X.JFP-18-073

24. Mieszkin, S., Furet, J.-P., Corthier, G., Gourmelon, M., 2009. Estimation of Pig Fecal Contamination in a River Catchment by Real-Time PCR Using Two Pig-Specific Bacteroidales 16S rRNA Genetic Markers. Appl. Environ. Microbiol. 75, 3045–3054. https://doi.org/10.1128/AEM.02343-08

25. Milbury, C.A., Zhong, Q., Lin, J., Williams, M., Olson, J., Link, D.R., Hutchison, B., 2014. Determining lower limits of detection of digital PCR assays for cancer-related gene mutations. Biomol. Detect. Quantif. 1, 8–22. https://doi.org/10.1016/j.bdq.2014.08.001

26. Mohan, S., Pascual-Garrigos, A., Brouwer, H., Pillai, D., Koziol, J., Ault, A., Schoonmaker, J., Johnson, T., Verma, M.S., 2021. Loop-Mediated Isothermal Amplification for the Detection of Pasteurella multocida, Mannheimia haemolytica, and Histophilus somni in Bovine Nasal Samples. ACS Agric. Sci. Technol. 1, 100–108. https://doi.org/10.1021/acsagscitech.0c00072

27. Nshimyimana, J.P., Cruz, M.C., Thompson, R.J., Wuertz, S., 2017. Bacteroidales markers for microbial source tracking in Southeast Asia. Water Res. 118, 239–248. https://doi.org/10.1016/j.watres.2017.04.027

28. Ordaz, G., Angel Merino-Mascorro, J., Garcia, S., Heredia, N., 2019. Persistence of Bacteroidales and other fecal indicator bacteria on inanimated materials, melon and tomato at various storage conditions. Int. J. Food Microbiol. 299, 33–38. https://doi.org/10.1016/j.ijfoodmicro.2019.03.015

29. Pang, H., McEgan, R., Micallef, S.A., Pradhan, A.K., 2018. Evaluation of meteorological factors associated with pre-harvest contamination risk of generic Escherichia coli in a mixed produce and dairy farm. Food Control 85, 135–143. https://doi.org/10.1016/j.foodcont.2017.08.003

30. Pascual-Garrigos, A., Maruthamuthu, M.K., Ault, A., Davidson, J.L., Rudakov, G., Pillai, D., Koziol, J., Schoonmaker, J.P., Johnson, T., Verma, M.S., 2021. On-farm colorimetric detection of Pasteurella multocida, Mannheimia haemolytica, and Histophilus somni in crude bovine nasal samples. Vet. Res. 52, 126. https://doi.org/10.1186/s13567-021-00997-9

31. Pendergraph, D.P., Ranieri, J., Ermatinger, L., Baumann, A., Metcalf, A.L., DeLuca, T.H., Church, M.J., 2021. Differentiating Sources of Fecal Contamination to Wilderness Waters Using Droplet Digital PCR and Fecal Indicator Bacteria Methods. Wilderness Environ. Med. 32, 332–339. https://doi.org/10.1016/j.wem.2021.04.007

32. Perkins, G., Lu, H., Garlan, F., Taly, V., 2017. Chapter Three - Droplet-Based Digital PCR: Application in Cancer Research, in: Makowski, G.S. (Ed.), Advances in Clinical Chemistry. Elsevier, pp. 43–91. https://doi.org/10.1016/bs.acc.2016.10.001

33. Ranjbaran, M., Verma, M.S., 2022. Microfluidics at the interface of bacteria and fresh produce. Trends Food Sci. Technol. 128, 102–117. https://doi.org/10.1016/j.tifs.2022.07.014

34. Ravaliya, K., Gentry-Shields, J., Garcia, S., Heredia, N., Fabiszewski de Aceituno, A., Bartz, F.E., Leon, J.S., Jaykus, L.-A., 2014. Use of Bacteroidales Microbial Source Tracking To Monitor Fecal Contamination in Fresh Produce Production. Appl. Environ. Microbiol. 80, 612–617. https://doi.org/10.1128/AEM.02891-13

35. Rumky, J., Kruglova, A., Repo, E., 2022. Fate of antibiotic resistance genes (ARGs) in wastewater treatment plant: Preliminary study on identification before and after ultrasonication. Environ. Res. 215, 114281. https://doi.org/10.1016/j.envres.2022.114281

36. Shanks, O.C., Atikovic, E., Blackwood, A.D., Lu, J., Noble, R.T., Domingo, J.S., Seifring, S., Sivaganesan, M., Haugland, R.A., 2008. Quantitative PCR for Detection and Enumeration of Genetic Markers of Bovine Fecal Pollution. Appl. Environ. Microbiol. 74, 745–752. https://doi.org/10.1128/AEM.01843-07

37. Siefring, S., Varma, M., Atikovic, E., Wymer, L., Haugland, R.A., 2008. Improved real-time PCR assays for the detection of fecal indicator bacteria in surface waters with different instrument and reagent systems. J. Water Health 6, 225–237. https://doi.org/10.2166/wh.2008.022

38. Silkie, S.S., Nelson, K.L., 2009. Concentrations of host-specific and generic fecal markers measured by quantitative PCR in raw sewage and fresh animal feces. Water Res., Cross-validation of detection methods for pathogens and fecal indicators 43, 4860–4871. https://doi.org/10.1016/j.watres.2009.08.017

39. Strawn, L.K., Fortes, E.D., Bihn, E.A., Nightingale, K.K., Gröhn, Y.T., Worobo, R.W., Wiedmann, M., Bergholz, P.W., 2013a. Landscape and Meteorological Factors Affecting Prevalence of Three Food-Borne Pathogens in Fruit and Vegetable Farms. Appl. Environ. Microbiol. 79, 588–600. https://doi.org/10.1128/AEM.02491-12

40. Strawn, L.K., Gröhn, Y.T., Warchocki, S., Worobo, R.W., Bihn, E.A., Wiedmann, M., 2013b. Risk Factors Associated with Salmonella and Listeria monocytogenes Contamination of Produce Fields. Appl. Environ. Microbiol. 79, 7618–7627. https://doi.org/10.1128/AEM.02831-13

41. Wang, J., Davidson, J.L., Kaur, S., Dextre, A.A., Ranjbaran, M., Kamel, M.S., Athalye, S.M., Verma, M.S., 2022. Paper-Based Biosensors for the Detection of Nucleic Acids from Pathogens. Biosensors 12, 1094. https://doi.org/10.3390/bios12121094

42. Wang, J., Dextre, A., Pascual-Garrigos, A., Davidson, J.L., Maruthamuthu, M.K., McChesney, D., Seville, J., Verma, M.S., 2021. Fabrication of a paper-based colorimetric molecular test for SARS-CoV-2. MethodsX 8, 101586. https://doi.org/10.1016/j.mex.2021.101586

43. Wang, J., Ranjbaran, M., Ault, A., Verma, M.S., 2023. A loop-mediated isothermal amplification assay to detect Bacteroidales and assess risk of fecal contamination. Food Microbiol. 110, 104173. https://doi.org/10.1016/j.fm.2022.104173

