## Supporting Information for "*Bacteroidales* as a Fecal Contamination Indicator in Fresh Produce Industry: A Baseline Measurement"

### for

**Supporting Figure**

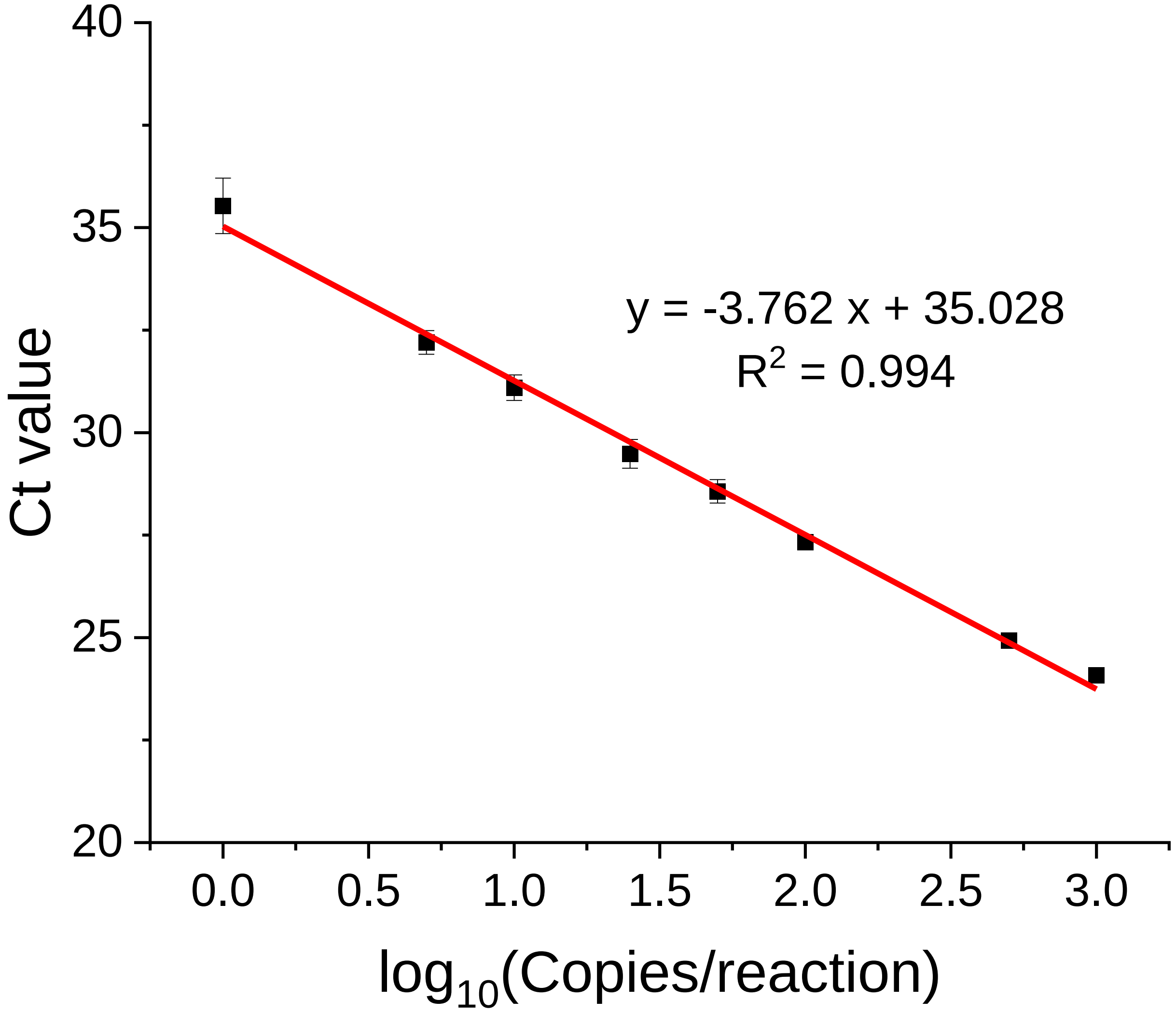


Figure S1: qPCR calibration curve. Serially diluted (1000 copies/reaction to 1 copy/reaction) *B. fragilis* genomic DNA (1 µL) samples were added to reactions (19 µL reagents) in triplicates. For NTC, 1 µL of nuclease-free water was added to the reaction mix instead of genomic DNA. The final volume of the reactions was 20 µL, and master mix used was NEB 2X Luna® Universal Probe qPCR Master Mix. The Ct values were calculated using software qPCRsoft 4.1 (baseline correction: 5, auto threshold) (Analytik Jena, Germany).
